# Novel Alzheimer risk genes determine the microglia response to amyloid-β but not to TAU pathology

**DOI:** 10.1101/491902

**Authors:** Annerieke Sierksma, Ashley Lu, Evgenia Salta, Renzo Mancuso, Jesus Zoco, David Blum, Prof Luc Buée, Prof Bart De Strooper, Mark Fiers

## Abstract

**Background:** Thousands of SNPs associated with risk of Alzheimer’s disease (AD) in genome-wide association studies (GWAS) do not reach genome-wide significance. When combined, they contribute however to a highly predictive polygenic risk score. The relevance of these subthreshold risk genes to disease, and how their combined predictive power translates into functionally relevant disease pathways, is unknown. We investigate here at the genome-wide level and in an unbiased way to what extent AD risk genes show altered gene expression in the context of increasing Aβ or Tau pathology in mouse models of AD.

**Methods:** We used an existing GWAS data set to generate lists of candidate AD genes at different levels of significance. We performed transcriptomic analysis on wild-type and transgenic APP/PS1 (APPtg) and Thy-TAU22 (TAUtg) mouse models at early and late stage of disease. We used unbiased weighted gene co-expression network analysis (WGCNA) to identify clusters of co-regulated genes responsive to Aβ or TAU pathology. Gene set enrichment was used to identify clusters that were enriched for AD risk genes.

**Findings:** Consistent and significant enrichment of AD risk genes was found in only one out of 63 co-expression modules. This module is highly responsive to Aβ but not to TAU pathology. We identify in this module 18 AD risk genes (p-value=9.0e-11) including 11 new ones, *GPC2, TREML2, SYK, GRN, SLC2A5, SAMSN1, PYDC1, HEXB, RRBP1, LYN* and *BLNK.* All are expressed in microglia, have a binding site for the transcription factor *SPI1* (PU.1), and become significantly upregulated when exposed to Aβ. A subset regulates FC-gamma receptor mediated phagocytosis.

**Interpretation:** Genetic risk of AD is functionally translated into a microglia pathway responsive to Aβ pathology. This insight integrates aspects of the amyloid hypothesis with genetic risk associated to sporadic AD.

## Introduction

Genetic background strongly determines the risk of sporadic Alzheimer’s Disease (AD).^1^ Apart from the APOE4 polymorphism and 42 other genetic loci, thousands of SNPs associated with risk of AD do not reach genome-wide significance.^2^–^4^ Polygenic risk scores (PRSs) incorporate the contribution of these variations and relate that to disease risk.^5^ PRSs for AD currently reach a prediction accuracy of 84%, albeit that a major proportion can be attributed to APOE status alone.^6^ A crucial question is whether AD risk genes functionally link to amyloid-β (Aβ) or TAU pathology or whether they define many parallel pathways that all lead to AD. We expect that at least part of the genes and pathways implicated in genome-wide association studies (GWAS) affect the cellular response of the brain to Aβ or TAU pathology.^7^ Such a model integrates parts of the amyloid hypothesis with the complex genetics of AD, which will lead to a more coherent view on the pathogenesis of AD.

Profiling of postmortem brain tissue only provides insights into the advanced stages of AD and cannot delineate cause-consequence relationships, which is required to develop mechanistic models for the pathogenesis of AD. Transgenic mouse models only partially recapitulate AD or frontotemporal dementia (FTD) phenotypes, but they provide detailed functional insights into the initial steps of disease, which is of high relevance for preventative therapeutic interventions.^8^ What is lacking until now, however, is the integration of functional information from mouse studies with the GWAS data obtained in human. By doing so, it might be possible to determine whether sub-significant AD risk genes, are involved in the cellular response to Aβ or TAU pathology. This would increase confidence that these genes are truly involved in AD and indicate in which pathways these genes play a functional role.

Here, we perform transcriptional profiling of mouse hippocampus after exposure to Aβ or TAU pathology, at early (4 months of age (4M)) and mature stages of disease (10M). We used APP^swe^/PS1^L166P^ (APPtg) and Thy-TAU22 (TAUtg) mice, both expressing the transgene from a *Thy1.2* promotor.^9,10^ Despite similar robust cognitive phenotypes, APPtg mice develop severe age-dependent transcriptional deregulation, while TAUtg mice have a milder and over time more stable molecular phenotype. AD risk genes uniquely converge in APPtg mice into a coordinated deregulated multicellular gene network that is strongly enriched in neuroinflammatory functions. Our work provides evidence that a large part of the genetic risk of AD is determining the microglial response to Aβ, and promotes 11 candidate GWAS genes for future AD research.

## Results

At 4M of age, APPtg and TAUtg mice are cognitively intact with mild levels of pathology, whereas they display at 10M overlapping profiles of hippocampus-dependent mnemonic deficits and substantial pathology.^9–11^ mRNAseq was performed on the hippocampus of 4M and 10M APPtg and TAUtg mice (TG) and their respective wild-type (WT) littermates, with n=12 per group and n=96 in total, yielding on average 7.7 million reads per sample (see Fig.1A).

**Fig. 1:**
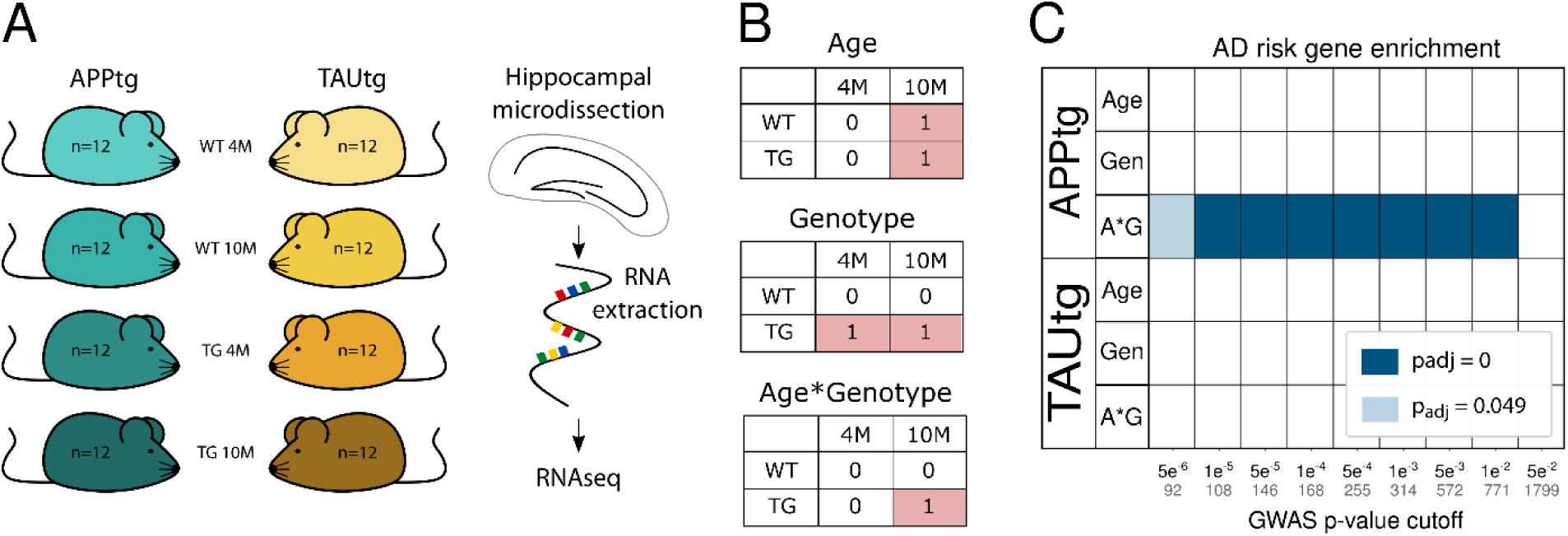
Enrichment of AD risk genes in APPtg and not in TAUtg mice. A) Experimental design for mRNA sequencing using n=12 per experimental group. B) Explanation of the 2×2 linear model, where those cells labeled with 1 are compared to the cells labeled with 0. In the age comparison, mRNA expression in all 10 month old (10M) mice is compared to all 4 month old (4M) mice. In the genotype comparison, mRNA expression in all transgenic (TG) mice is compared to all wild-type (WT) mice. In the age*genotype comparison, we assess which transcripts are differentially expressed in the 10M TG mice compared to all other groups. C) Based on Marioni et al. ^2^ various sets of AD GWAS risk genes were created using different cut-off p-values indicated on the x-axis (number of genes within each set is written in grey). Enrichment for AD risk genes was assessed among the different statistical comparisons for APPtg and TAUtg mice (Int: age*genotype, Gen: genotype). Colors represent –log10(Benjamini-Yuketieli adjusted p-value) for the enrichment; blank means no significant enrichment.

### Enrichment of AD risk genes is only found in the transcriptomic response of APPtg mice

A 2×2 linear model (Fig.1B) was employed to investigate the effects of genotype, age, and age*genotype interaction. The age comparison identifies transcripts changed between 4M and 10M old mice (Fig.2A). The genotype comparison shows mRNAs different between WT and TG mice (Fig.2B). The age*genotype interaction, finally, assesses which transcripts change with aging uniquely in the TG mice (Fig.2C). The study thus reflects the transcriptional changes manifesting in the mice at two critical time points: initially when the first signs of Aβ and TAU pathology occur and later on, when the biochemical alterations are manifest and accompanied by cognitive deficits.

We wondered whether GWAS-based AD risk genes would be equally responsive to Aβ or TAU pathology. We included both established AD risk genes, i.e. genes with p-value<5×10e-8 in various GWAS studies, as well as subthreshold AD risk genes as these contribute significantly to AD risk predictions through polygenic inheritance.^6^ We examined multiple sets of such genes taken from Marioni et al., which combines UK Biobank AD-by-proxy data with the IGAP database.^2^ By using arbitrary p-value cut-offs with decreasing significance for AD association, genes sets of increasing size were created (see Fig.1C and Supplementary Table 2). PRS studies have demonstrated that GWAS SNPs up to a cutoff of p<0.5 still improve the predictive power of risk score. We decided to limit our study to a cutoff of p<0.05, yielding an already large set of 1799 genes. The enrichment of these AD risk gene sets in the different transcriptional responses of the mice was assessed using gene set enrichment analysis (GSEA). The data (Fig.1C) demonstrate that independently of the set size, ranging from 92 genes (p<5e-6) to 1799 genes (p<5e-2), AD risk genes are found consistently, and significantly (padj<1e-250) enriched among the genes changing as APPtg mice age (“APP interaction” Fig.1C), but not in Tautg mice. The smallest set (n=92 genes with p<5e-06), also significantly enriches among the APPwt vs APPtg comparison (“APP genotype”, Fig.1C; padj=0.0057). This gene set contains many microglia-expressed genes e.g. *Treml2, Inpp5d* or *Gal3st4,* (see Fig2D and Supplementary Table 1). Thus, genes that enhance the risk of AD are clustering among genes that are deregulated over time with increasing Aβ-but not TAU-pathology.

**Fig. 2:**
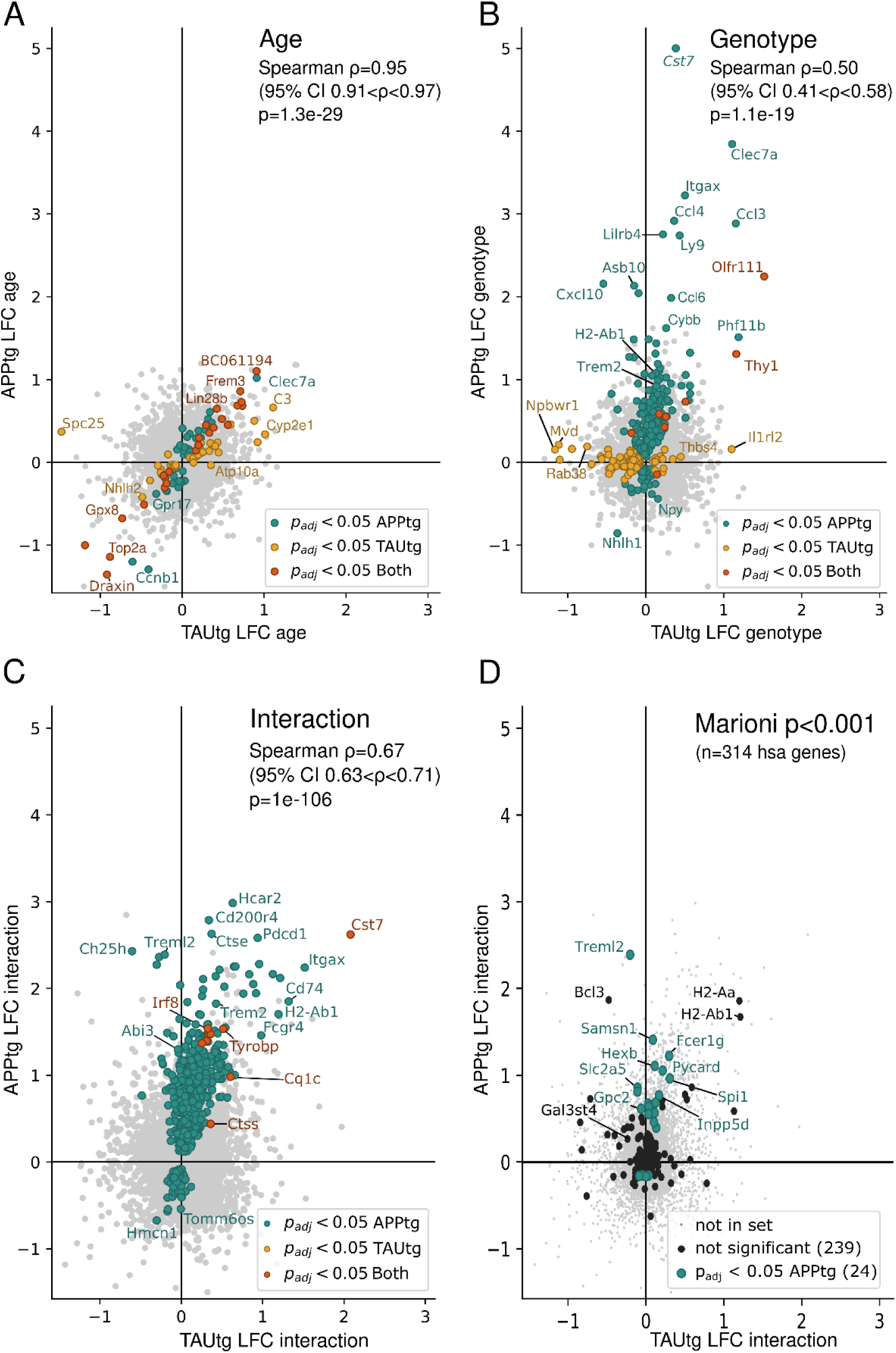
Changes in gene expression exacerbate with aging in APPtg but not in TAUtg mice. Log2 fold change (LFC) in TAUtg (x-axis) and APPtg mice (y-axis) after differential expression analysis, assessing the effects of age (A), genotype (B) and age*genotype interaction and where in D only the Marioni-based GWAS genes at p<0.001 are depicted. Upregulated genes are on the right part (TAUtg mice) or upper part (APPtg mice) of the graph; downregulated genes are on the left part (TAUtg) or lower part (APPtg) of the graph. Colored dots represent significantly differentially expressed genes (Benjamini-Yuketieli-adjusted p-values (Padj) <0.05) for APPtg (green dots), TAUtg (yellow dots) or for both (red dots). Spearman correlation assesses the correlation between APPtg and TAUtg mice when ranking genes that are significantly differentially expressed in either APPtg or TAUtg mice from most up-to most down-regulated on a combined score of LFC and Padj (i.e. signed log10(p-value), where the sign is determined by the LFC). Black dots in D represent all GWAS genes with p<0.001.

### Changes in gene expression exacerbate with aging in APPtg but not in TAUtg mice

To assess the functional substrate of the AD risk gene enrichment in APPTg mice, we compared the transcriptional deregulation in the two mouse models in more detail (see Fig.2A-C and Supplementary Table 1). The transcriptional response of the APPtg and TAUtg mice caused by aging (i.e. independent of transgene) is practically identical (Spearman correlation R=+0.95, p=1.3e-29, 95% confidence interval (CI) +0.91 to +0.97; see Fig.2A). When comparing the effects of transgene expression only, the similarity between APPtg and TAUtg mice becomes rather moderate (R=+0.50, p=1.1e-19, 95% CI=+0.41 to +0.58; see Fig.2B) and this is only slightly enhanced in the interaction model of age*genotype (R=+0.67, p=1e-106, 95% CI=+0.63 to +0.71; see Fig.2C). Thus, while both mouse models age in similar ways, major differences in the transcriptional response between APPtg and TAUtg mice show that these are very different pathologies causing very divergent cellular reactions.

The *APP/PSEN1* transgene causes prominent changes (287 genes in total) in gene expression (green dots, Fig.2B) with most (n=219, i.e. 76%) genes upregulated (log2-fold change (LFC):+0.07 to +5.00, Benjamini-Yekutieli adjusted p-value (padj)<0.05). When the aging component is added (i.e age*genotype) even more genes become upregulated (623 mRNAs (i.e.78%), LFC:+0.12 to +2.98, padj<0.05), while also 175 genes down-regulate their expression (LFC:-0.67 to −0.08, padj<0.05) (Fig.2C). The many up-regulated genes in APPtg are often involved in microglia (Supplementary Fig.S2) and neuroinflammatory responses, including *Tyrobp* (LFC genotype (G):+1.19, LFC age*genotype (A*G):+1.53), *Cst7* (LFC G:+5.00, LFC A*G:+2.62) and *Itgax* (LFC G:+3.22, LFC A*G:+2.24). These changes are strong, up to 32-fold. The enriched AD risk genes (Fig.1C) are predominantly upregulated in the age*genotype comparison (Fig.2D). Thus, it appears that an increased expression of neuroinflammatory genes, most likely of microglial source, is an early (Fig.2B) and persistent (Fig.2C) component of Aβ pathology and many AD risk genes follow a similar transcriptional response.

TAUtg mice show markedly fewer transcriptional changes with very little aggravation over time. In the genotype comparison (TAUwt vs TAUtg), only 47 genes become significantly up-regulated (LFC:+0.06 to +1.52; padj<0.05) and 77 down-regulated (LFC:-1.30 to −0.05, padj<0.05; Fig.2B, yellow dots). Only 9 genes are up-regulated (LFC:+0.25 to +0.60, padj<0.05; Fig.2C, red dots) when genotype is combined with age in the interaction model. The majority of deregulated genes (62%) in TAUtg vs TAUwt show decreased expression (Fig.2B) and are of neuronal origin (Supplementary Fig.S2). The 9 up-regulated genes in the age*genotype comparison of TAUtg mice are overlapping with APPtg mice and are of microglial (*C1qa, C1qc, Tyrobp, Ctss, Irf8, Mpeg1, Cst7, Rab3il1*) or astroglial (*Gfap*) origin. The overlap with APPtg reflects a (much milder) microglia response in Tautg. With the exception of Cst7 (LFC: 2.08), the upregulation is indeed very modest (average LFC of 8 others: 0.38) compared to APPtg mice (max LFC: 2.98; average LFC: 0.70). Overall, we can conclude that the molecular, pathobiological and cellular responses in APPtg and TAUtg appear fundamentally different. APPtg drives a strong, over time exacerbating inflammatory response, while the Tau transgene causes an early effect on gene expression of genes related to neuronal functions. Remarkably, while the underlying pathogenic mechanisms in the two models are very different, they converge into a very similar and robust cognitive phenotype. Most importantly, genes associated with genome wide statistical significance to AD as well as genes below that threshold, are transcriptionally active when facing accumulating Aβ, but not TAU pathology.

### AD risk genes are co-regulated in a specific functional gene expression module

Next, we performed unbiased weighted gene co-expression network analysis (WGCNA) on each mouse model separately to investigate whether AD risk genes would cluster in functional modules. We obtained in total 63 modules (Supplementary Fig S3+4). GSEA with the GWAS gene set generated from Marioni et al.^2^, at different cut-offs for statistical significance as explained above (Fig.3A and Supplementary Table 2) demonstrated that the largest set of risk genes (e.g. n=1799 genes at p<0.05), enrich among 4 APPtg-and TAUtg-based modules (Turquoise, Blue, Fig.3A and Supplementary Fig.S3 and S4). However, when taking gene sets defined by increasing statistical significance (p<0.001 or smaller), the only module that persistently demonstrates significant enrichment with GWAS genes is the APPtg-Blue module (see Fig.3A). The APPtg-Blue module is large (n=4236), and contains 62% of all the genes significantly differentially expressed in aging APPtg mice (age*genotype, which is more than expected by chance (log2 odds ratio (LOR): 2.90, p=1.54e-158)). We assume that this module provides the integrated and coordinated response of the brain to amyloid pathology. It is important to stress that despite the module being large, still significantly more AD GWAS genes are in the APPTg-Blue module than expected by chance alone (1e-250<padj<0.01). Thus, it appears that genes associated with increasing risk of AD, generally cluster within this module.

**Fig. 3:**
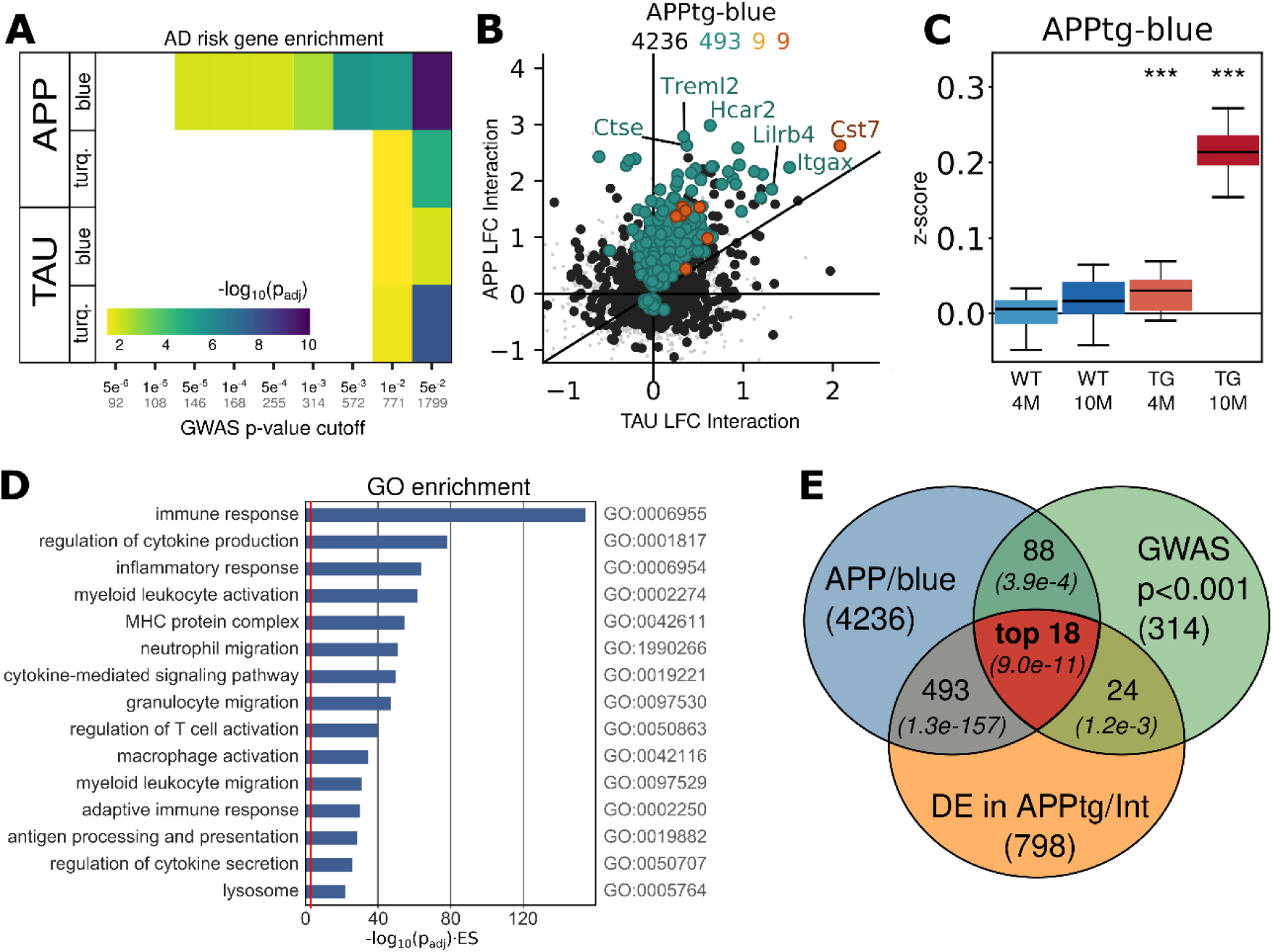
The APPtg-Blue module represents a coordinated transcriptional response only present in APPtg mice. A) Gene set enrichment analysis using Fisher’s exact test of GWAS genes from Marioni et al. ^2^ at different p-value cut-offs among the WGCNA-derived modules in APPtg or TAUtg mice (see also Supplementary Fig.S3+4). Colors represent –log10(Benjamini-Yuketieli adjusted p-value) for the enrichment. Numbers on the x-axis: black = p-value cut-off; grey = size of GWAS gene set. B) Log2 fold change (LFC) in TAUtg (x-axis) and APPtg mice (y-axis) after differential expression analysis, assessing the effects of age*genotype interaction. Color code for dots/numbers: grey = genes in hippocampus (n=15824); black = genes in APPtg-Blue (n=4236); green = genes significantly differentially expressed in APPtg mice (n=493); yellow = significantly differentially expressed genes in TAUtg mice (n=9); red = significantly differentially expressed genes in both (n=9). C) Z-score distribution per experimental group for all genes within the APPtg-Blue module. Boxplots: center line, median; box limits, 25^th^-75^th^ quartiles; whiskers, 1.5x interquartile range. Empirical p-values are Bonferroni adjusted (pbonf<0.001) and indicate significant shift in z-score distribution (see Supplementary Materials & Methods). D) Gene Ontology (GO) enrichment for genes within the APPtg-Blue module. The x-axis depicts –log10(FDR-adjusted p-value) multiplied by the enrichment score (ES), where the red line represents an – log10(0.049)*(ES=1). E) The “top 18” GWAS genes are prioritized by finding the intersection of genes within the APPtg-Blue module (n=4236), AD GWAS genes with p<0.001 in Marioni et al. ^2^ (n=314) and significantly differentially expressed genes in the age*genotype comparison of APPtg mice (n=798).

Remarkable, most of the differentially expressed genes in this module are upregulated (Fig.3B+C). We functionally characterized this Aβ-induced transcriptional response. The APPtg-Blue module shows a highly significant overlap with the microglia-specific gene set (LOR: 1.90, padj=1.74e-77; Fig.4B and Supplementary Table 3) and to a lesser extent the astrocyte gene set (LOR: 0.54, padj: 0.014). Moreover, this module is highly enriched for GO categories involving immune response, cytokine production and inflammation (Fig.3D and Supplementary Table 4). It furthermore shows a highly significant overlap with a recently published Aβ-response network (LOR: 2.0, p=2.2e-16) as well as with the microglia-immune module derived from the brains of late-onset AD patients (LOR: 2.14, p=8.0e-73), demonstrating that this transcriptional response to increasing Aβ is similar in this Aβ mouse model and AD patients.^12,13^

**Fig. 4:**
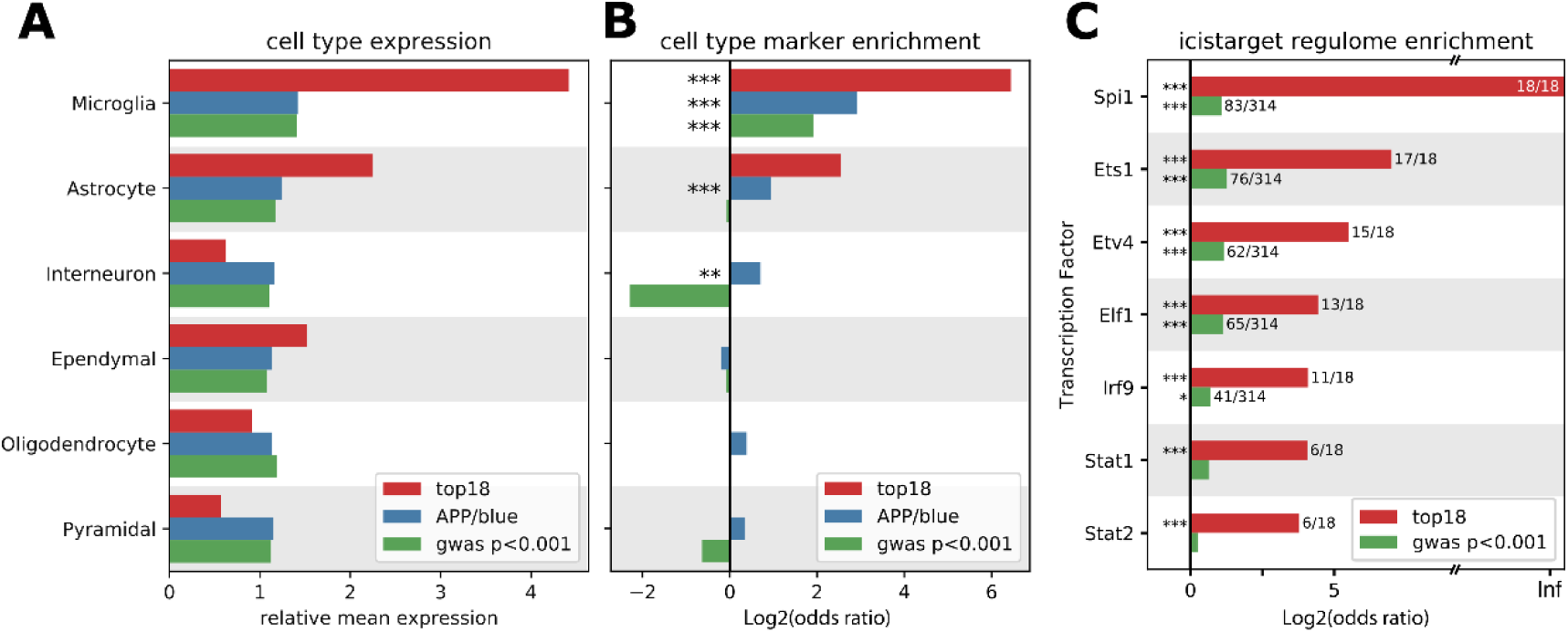
“Top 18” genes are expressed in microglia and regulated by *Spi1*. A, B) Average expression of genes within all cell types (A) and enrichment of the cell type marker genes as determined by Zeisel et al. ^19^ (B) among the ‘top 18’ genes (red bars) as defined in Fig.3E, the genes within the APPtg-Blue module (blue bars) and GWAS genes with p<0.001 (^2^; green bars). C) Enrichment of transcription factor targets among the ‘top 18’ genes (red bars) and the GWAS genes with p<0.001 (green bars). ***: significant enrichment, p< 0.001; **: p < 0.01. *: p<0.05. P-values are Benjamini-Yuketieli-adjusted, except for (C) where they are empirical and Bonferroni-adjusted.

We identified transcription factors potentially regulating this module. Out of 16 APPtg-Blue-associated transcription factors, *Spi1* (a.k.a. PU.1) comes out as the top candidate, which is particularly interesting as a SNP within *Spi1* is protective in AD.^14^ *Spi1* is also significantly differentially expressed in the APPtg age*genotype interaction comparison (LFC: 0.96, padj=9.92e-05). Other transcription factors predicted to regulate the APPtg-Blue module include microglia-related and interferon-responsive transcription factors Stat2, Stat1, Ets1 and Irf7, although only Stat1 was significantly differentially expressed in the APPtg age*genotype comparison (LFC: 0.39, padj=0.0013). To summarize, we can conclude that the APPtg-Blue module shows a coordinated transcriptional response to increasing Aβ load employing a large number of AD risk genes. This cellular response involves microglial and astrocyte genes and seems, at least partially, regulated by the transcription factor *Spi1*.

Of note, no such module enriched for AD risk genes is found in the TAUtg mice (see Fig.3A). Several smaller modules show significant overlaps with the microglial gene set (TAU-Red LOR: 2.06, padj=5.33e-35; TAU-Paleturquoise LOR: 2.14, padj=0.013), the astrocyte gene set (TAU-Red LOR: 1.10, padj: 3.8e-03), and the pyramidal neuron gene set (e.g. TAU-Black LOR:1.28, padj:7.64e-12 and TAU-Pink LOR:1.02, padj: 3.6e-03; see Supplementary Fig.S3+4 and Supplementary Table 3), suggesting that the response to Tau pathology is also multicellular by nature. However, the fragmented cellular response to TAU pathology is clearly different from the coordinated transcriptional regulation response in the APPtg-Blue module.

### Prioritization of subthreshold GWAS risk genes of AD within the Aβ-induced transcriptional response

The set of 1799 GWAS genes (p<0.05) contains 439 genes co-regulated within the APPtg-Blue module, further demonstrating that this constitutes a core pathobiological response to Aβ onto which genetic risk for sporadic AD converges. These 439 risk genes encompasses well established (“canonical”) AD genes, i.e. *Abi3, Apoe, Bin1, Clu, Def8, Epha1, Fcer1g, Fermt2, H2-Ab1* (*HLA-DQA* in humans), *Inpp5d, Plcg2, Prss36, Rin3, Siglech* (*CD33* in humans), *Spi1, Tomm40, Trem2* and *Zcwpw1.* The 421 other genes in this data set have been associated to AD with decreasing degrees of statistical certainty. When taking gene sets with increasingly stringent p-values for their association to AD risk, the number of GWAS genes in the selection obviously drops (Supplementary Table 2), but the genetic evidence for the role of these genes in AD increases. In the end, the choice of the p-value threshold is arbitrary, as even SNPs with p-values up to 0.5 contribute to the PRS and therefore contain information.^6^

The important insight is, however, that identified genes are implied in AD by two <u>independent</u> unbiased experimental approaches here: genetically, by contributing to the risk of AD in human studies, and functionally, by being part of the APPtg-Blue-response module in a mouse model for AD.^2,6^ We focus here on GWAS genes that: 1) have been associated to AD with p<0.001 as these genes are more likely to make an individual impact on disease, 2) that are present in the APPtg-Blue module, and 3) that are, additionally, significantly differentially expressed in the age*genotype comparison of APPtg mice (see Fig.3E). The latter criterion is restrictive because we require that the gene is not only part of the APPtg-Blue response module but is also significantly altered in expression, thus part of the core response that defines the module. These three criteria show a highly significant intersection of 18 genes which is markedly better than expected by chance alone (expected 2.2 genes, padj=9.0e-11, SuperExactTest ^15^, Fig.3E and Supplementary Table 7). These 18 genes are *APOE, CLU, GPC2, INPP5D, CD33, PLCG2, TREML2, SPI1, FCER1G, SYK, GRN, SLC2A5, SAMSN1, PYDC1, HEXB, RRBP1, LYN* and BLNK (see Fig.5). The full set of GWAS genes with p<0.5 and significantly differentially expressed in the APPtg-Blue module (n=263 genes) are listed in Supplementary Table 6.

**Fig. 5:**
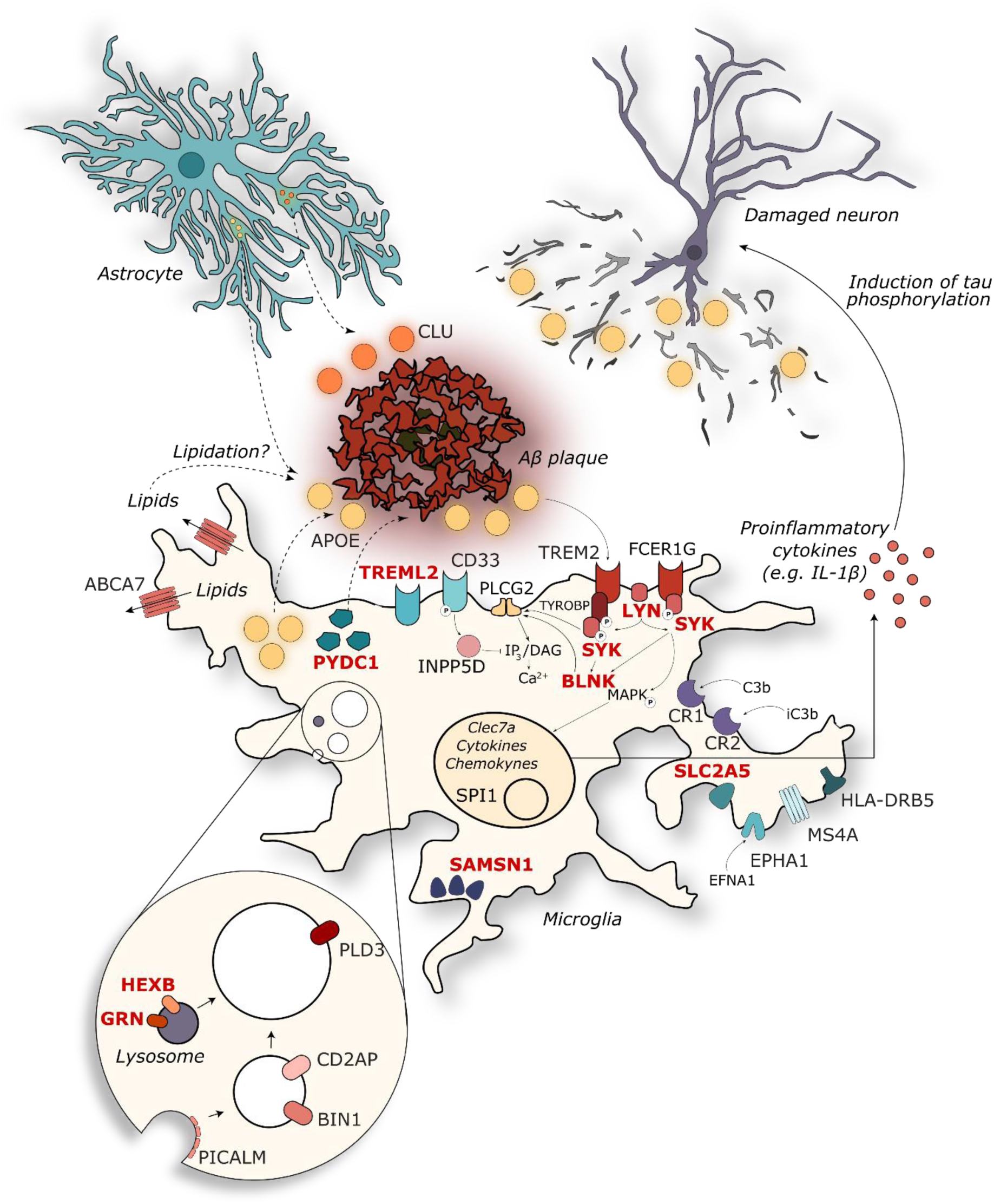
The 11 novel prioritized AD risk genes mainly cluster together with other established AD genes within microglial or lysosomal gene networks involving innate immunity. Genes are depicted in their known cellular components, with previously described established GWAS ^3,4^ in black, and the prioritized novel genes (this paper) in red. See discussion section for further functional annotation and references.

Among the 18 genes we find 7 established AD GWAS genes, i.e. reaching genome wide significance in one or more studies.^3,4,16^ We indicate them here together with the p-value (*p*_*mar*_) for the GWAS association as listed in Marioni et al. ^2^: *APOE* (p_mar_: 2.80e-118), *CLU* (p_mar_: 7.68e-15), *INPP5D* aka SHIP1 (p_mar_: 2.42e-10), *CD33* (Siglech in mice; p_mar_: 1.82e-07), *PLCG2* (p_mar_: 2.76e-06), *SPI1* (p_mar_: 5.45e-06) and *FCER1G* (p_mar_:1.10e-05). The other 11 genes have not reached genome-wide statistical significance in any previously published GWAS study but are coming forward here based on additional significant observations, i.e. that these genes are significantly differentially expressed when exposed to pathology and are present in the APPtg-Blue module as discussed. Although each of these observations have an associated p-value, they are derived from different hypotheses, making it difficult to combine them into a specific p-value per gene. However, by using an extension on the classical Fischer’s Exact test (SuperExactTest ^15^) to evaluate the significance of finding overlaps between multiple gene sets, it was demonstrated that observing an intersection of 18 genes among these three gene sets is highly significantly better than expected by chance ((padj = 9.0e-11, see Supplementary Table 7 and Fig.3E). Among these 11 novel genes (their individual p_mar_ is again indicated), *SYK* (p_mar_: 1.41e-04), *LYN* (p_mar_: 8.38e-04) and *BLNK* (p_mar_: 9.52e-04) together with the already mentioned ‘canonical’ genes *INPP5D, FCER1G* and *PLCG2*, play a role in FC-gamma receptor mediated phagocytosis (see also Fig.5). When examining the longer list of priority GWAS genes (see Supplementary Table 6), we can find more members of this pathway, including *TREM2, ITGAM* and *VAV1*. Moreover, the same genes, as well as others of the top 18 list, i.e. *PYDC1/PYCARD* (p_mar_: 4.69e-04), *GRN* (p_mar_: 1.56e-04), *SLC2A5* (p_mar_: 2.07e-04), and *SAMSN1* (p_mar_: 4.66e-04) are all part of the ‘immune and microglia’ module regulated by *TYROBP* inferred by Zhang et al. from RNAseq data derived from late-onset AD patients.^13^ Other microglia-and lysosome-related genes are *TREML2* (p_mar_: 4.90e-06) and *HEXB* (p_mar_: 4.87e-04). *RRBP1* (p_mar_: 7.10e-04) is an ER-associated ribosomal protein ^17^, with high expression in glial cells (Supplementary Table 6). All of the 18 AD risk genes are predicted *SPI1* targets according to i-cisTarget (see Fig.4C) and 11 out of these (*APOE, BLNK, HEXB, INPP5D, LYN, PLCG2, RRBP1, SAMSN1, SLC2A5, SPI1, SYK)* are demonstrated targets in a ChIPseq experiment in the BV2 microglia cell line.^18^ Thus many of these subthreshold AD risk genes that contribute to polygenic inheritance, participate in microglia-related functional networks both in AD patients as well as in mouse models of Aβ pathology. We calculated the average expression of each gene within each cell type for the top 18 genes, using the data compiled by Zeisel et al., which demonstrated that the top 18 genes are highly expressed in microglia (~4 times more than average, see Fig.4A).^19^ Thus, these top 18 GWAS genes contribute to the cellular response of microglia when facing increasing Aβ levels. The possible role of these genes together with other microglia-related previously established GWAS genes is schematically summarized in Fig.5, and further discussed below.

## Discussion

This study identifies a gene transcription module (APPtg-Blue) that is specifically induced by overexpression of mutated *APP/PSEN1* (but not by *MAPT*) and is enriched for a large set of genes that were previously linked with SNPs associated to AD risk without reaching genome-wide significance. Our experiments suggest that these genes are part of a pathway that characterizes the microglia response to Aβ (see Fig.5). While the individual effects of various SNPs on expression of these genes are likely small and biological insignificant, the combination of hundreds of such subtle alterations, if synergic in one pathway, might veer the cellular response to Aβ in a disease causing direction through biological epistasis.^20^

Functionally, this co-regulatory network (the APPtg-Blue module), represents a large neuroinflammatory response involving astroglia and, overwhelmingly, microglia. These observations considerably expand previous work, strengthening the hypothesis that the major response in brain to Aβ pathology is neuroinflammation.^21,22^ Our and other data suggest that an early and profound neuroinflammatory response is an integral and perhaps even driving component in AD pathogenesis, and that Aβ is sufficient to induce this innate immune reaction.^13,22^ Our data strengthen this concept considerably by demonstrating that a large part of the GWAS-based genetic risk affects genes that are expressed in the molecular pathway defining this neuroinflammatory response to Aβ pathology.

Earlier studies have relied heavily on functional analysis and module connectivity for putative gene selection involved in AD pathogenesis, but whether these genes were also differentially expressed was not considered.^12,13,22^ The 5 recently reported novel AD risk genes *OAS1, ITGAM, CXCL10, LAPTM5* and *LILRB4* are all found within the APPtg-Blue module, but only *Itgam* and *Laptm5* are significantly differentially expressed with increasing Aβ load.^12^ We used two independent functional criteria: genes must have a genetic association with AD at p<0.001 and show a statistically significant change in expression when exposed to increasing levels of Aβ over time, indicating that these genes are part of the cellular response. Therefore, we can predict with high confidence (BY-padj=9.0e-11) that they are indeed involved in an AD relevant disease pathway responding to Aβ pathology. It must be noted that the GWAS genes currently used were based on genomic proximity to AD-associated SNPs, which may result in spurious associations. Although this potential caveat may affect our outcome, it is unlikely that the expression of such an erroneously identified gene would be significantly altered in AD-relevant pathways and thus will not appear in our final list.

Among the prioritized SNPs, many are linked to known myeloid-expressed genes involved in innate immune pathways (see Fig.5). Briefly, *FCER1G* encodes for the gamma subunit of IgE-specific Fc receptors and plays a role in phagocytosis and microglial activation.^23^ *SYK* is recruited by *FCER1G* and *TYROBP* (DAP-12) upon activation of *TREM2*, whereas *LYN* is localized at the plasma membrane and mediates *TREM2*/*SYK* phosphorylation.^24,25^ *BLNK* is a target of *SYK* and mediates the recruitment of *PLCG2*.^25^ *TREML2*, unlike *TREM2*, is reported to activate microglia and promote release of pro-inflammatory cytokines.^26^ *PYCD1* (*Pycard* in mouse) encodes for apoptosis-associated speck-like protein containing C-terminal caspase recruitment domain (ASC), a core component of the inflammasome complex, and has been recently linked to seeding and spreading of Aβ in mouse models of AD.^27^ The lysosomal protein *HEXB* and glucose transporter *SLC2A5* are part of the microgliome profile.^28^ *GRN* can adopt multiple subcellular localizations from lysosomes (as granulin) to the extracellular space (as progranulin) and exerts a negative control upon microglial activation in mouse models (see Fig.5).^29^ Interestingly, all of the prioritized AD risk genes are *Spi1* targets, substantiating the pivotal role of this transcription factor in the observed Aβ-induced transcriptional response.^14,30^

It remains to be clarified where, when and how in this pathway TAU pathology starts and how it contributes to disease progression towards dementia, and whether Aβ directly, or indirectly by the broad and complex neuroinflammatory (APPtg-Blue) response, induces abnormal TAU phosphorylation or conformation. Our hypothesis implies, however, that TAU pathology comes downstream of the microglia/neuroinflammatory response on Aβ because GWAS-defined genetic risk for AD is not associated with TAU-induced gene expression changes. The TAUtg transcriptional analysis does suggests that TAU induces very early down-regulation of neuronal genes (Supplementary Fig.S2), which vibes with the observations that dementia correlates much better with the appearance of tangles than amyloid plaques. Besides its role in regulating immune response, the newly prioritized risk gene *Syk* is a known tau kinase, providing an interesting genetic link between Aβ and TAU pathology.^31^

In conclusion, we hypothesize that the transition from a rather asymptomatic biochemical or prodromal phase, to the clinical, symptomatic, phase in AD, is largely determined by the combined inheritance of low-penetrant SNPs, which, as our work suggests, might affect the gene expression in the APPtg-Blue module. The concept of PRS forces us to abandon the classical gene-based view on disease etiology and to consider GWAS genes as part of a functional network.^32^ Given the robustness of biological systems and (the theory of) genetic buffering, a single SNP within the network will not lead to disease.^33^ However, multiple SNPs within the same network may tip the balance to a disease-causing disturbance. Such hypothesis also provides an explanation for the conundrum that some patients with high Aβ burden succumb without clinical symptoms.^34^ While Aβ might be the trigger of the disease, it is the genetic make-up of the microglia (and possibly other cell types) which determines whether a pathological response is induced. Identifying which SNPs are crucial to such network disturbances and how those SNPs lead to altered gene expression will be the next big challenge.

## Materials & Methods

### Study design

The goal of the current study was to assess the enrichment of AD GWAS genes among the transcriptional responses as APPstg and TAUtg mice develop pathology over time. Given the mice’ age and genotype, mice could not be distributed randomly across the experimental groups, but randomization was performed during RNA extraction and library preparation to avoid batch effects. Library preparation was performed in a blinded fashion. Details regarding sample size, biological replicates and outliers can be found below.

### Mice

The local Ethical Committee of Laboratory Animals of the KU Leuven (governmental license LA1210591, ECD project number P202-2013) approved all animal experiments, following governmental and EU guidelines. APPtg (a.k.a. B6.Cg-Tg(Thy1-APPSw,Thy1-PSEN1*L166P)21Jckr)^9^) mice express *APP*^*Swe*^ and *PSEN1*^*L166P*^ transgenes, while TAUtg (a.k.a. THY-Tau22 or Tg(Thy1-MAPT)22Schd^10^) mice, express the 412 aa isoform of the human 4-repeat *MAPT* gene containing the G272V and P301S mutations. Both mice are made using the *Thy1.2* promoter. Littermate transgenic (TG) and wild-type (WT) male mice were sacrificed at 4 months (average 123.8 days, SD 1.84 days) or 10 months of age (average 299.8 days, SD 2.22 days), creating 8 experimental groups (n=12 per group; see Fig.1A). Number of mice was determined as described in Sierksma et al.^35^ Following cervical dislocation, hippocampi were microdissected, snap-frozen in liquid nitrogen and stored at −80°C.

### RNA extraction, library construction, sequencing and mapping

The left hippocampus of each mouse was homogenized in TRIzol (Invitrogen, Carlsbad, CA, USA) using 1ml syringes and 22G/26G needles and purified on mirVana spin columns according to the manufacturer’s instructions (Ambion, Austin, TX, USA). RNA purity (260/280 and 260/230 ratios) and integrity was assessed using Nanodrop ND-1000 (Nanodrop Technologies, Wilmington, DE, USA) and Agilent 2100 Bioanalyzer with High Sensitivity chips (Agilent Technologies, Inc., Santa Clara, CA, USA) and Qubit 3.0 Fluorometer (Life Technologies, Carlsbad, CA, USA), respectively. RNA integrity values of the samples ranged from 7.9 to 9.3 (median= 8.6).

Library preparation and sequencing was performed at the VIB Nucleomics Core (Leuven, Belgium) using 1 μg of total RNA per sample. Poly-A containing mRNA molecules were purified using Illumina TruSeq® Stranded mRNA Sample Prep Kit (protocol version 15031047 Rev.E) and poly-T oligo-attached magnetic beads. After reverse transcription with random primers, different barcodes were introduced to each sample by ligating a single ‘A’ base to the 3’ ends of the blunt-ended cDNA fragments and multiple indexing adapters and 10 cycles of PCR amplification were performed. 96 libraries were pooled and sequenced in 4 runs using NextSeq 500 High75 Output kits on an Illumina NextSeq 500 instrument (Illumina, San Diego, CA, USA). Reads were pre-processed and mapped using BaseSpace SecondaryAnalysis (version 2.4.19.6, basespace.illumina.com), filtering out abundant sequences and trimming 2bp from the 5’ end. Reads were aligned against the mm10/GRCm38 *Mus musculus* reference genome by Tophat2 (version 2.0.7, ^36^).

### Data pre-processing and differential expression analysis

We found in 2 of the 12 APPtg-10M mice a 46% lower expression levels of hsa-APPswe, a 26% lower expression of hsa-Psen1^L166p^ and a 24% reduction in mmu-Thy1 gene, which drives the expression of the transgenes. Principle component analysis (using log2(raw reads from Feature Counts +1) and the ‘prcomp’ function in R) also revealed that these 2 mice were closer to the 4M APPtg mice than the 10M APPtg mice (see Supplementary Fig.S1). In the absence of an explanation for this phenomenon, these two mice were excluded from all further analyses.

mRNAs with average raw read counts ≤ 5 in 10 out of 96 samples were discarded, leaving 15824 mRNAs for differential expression (DE) analysis. Non-biological variation due to library preparation technicalities (i.e. effects for library prep batch, RNA extraction group and RNA concentration) was removed by the *removeBatchEffect* function from limma package 3.22.7 Bioconductor/R.^37^ DE analysis was conducted using a 2-way interaction model (age, genotype, age*genotype, see Fig.1B; two-sided testing) for APPtg and TAUtg mice separately. To adjust for multiple testing, Benjamini-Yuketieli (BY) p-value adjustment was performed, as we want to control for false discovery rate across experiments that have partially dependent test statistics, hence the traditional Benjamini-Hochberg adjustment was not applicable.^38^ Further differential expression analyses were performed by comparing WT to TG mice at 4M and 10M separately, with BY p-value adjustment across all 4 comparisons. Ranking of genes for Spearman correlations (two-sided testing) was always performed on the basis of signed log10(p-value), the log10 of the unadjusted p-value with a positive sign if the LFC of that gene within that comparison was >0 and a negative sign if the LFC was <0.

### Cell-specific datasets

Cell type specific genes sets for pyramidal neurons (n=701 genes), interneurons (n=364), astrocytes (n=239), microglia (n=435), oligodendrocytes (n=452), endothelial (n=352) and ependymal cells (n=483) were derived from Zeisel et al. ^19^ (Supplementary Table 1). Using these gene sets and a count matrix that was z-score normalized across samples, we calculated for each cell type (*t*) the average z-score (*Z*_*tg*_) for each experimental group (*g*) and compared this to the respective 4M WT group. We assesses significance using empirically derived p-values (see Supplementary Materials and Methods and Supplementary Fig.S5).

### GWAS gene set enrichment analysis

Human GWAS genes derived from Marioni et al.^2^ were converted to mouse orthologues using the Ensemble Biomart Release 94.^39^ Using the Gene Set Enrichment Analysis Preranked module (Broad Institute ^40,41^, two-sided testing) enrichment for GWAS genes was tested among up-and downregulated genes sorted on signed log10(p-value) based on the different statistical comparisons (age, genotype and age*genotype interaction), for APPtg and TAUtg separately, and performing Benjamini-Yuketieli p-value adjustment.

### Weighted gene co-expression network analysis (WGCNA)

The WGCNA package in R was used to build unsigned mRNA coexpression networks for APPtg and TAUtg mice separately using all 15824 expressed genes.^42^ To generate an adjacency matrix with is the smallest threshold that satisfies the scale-free topology fit at R^2^=0.9, soft power 3 is used for APPtg and 4 for TAUtg mice, respectively. The topology overlap (TO) was calculated based on the adjacency matrix which measures the network interconnectedness. The topology overlap dissimilarity was then calculated by 1-TO and used as input for average linkage hierarchical clustering. Branches of the hierarchical clustering tree were then assigned into modules using cutreeHybrid from the dynamicTreeCut package (deepSplit = 2, minModuleSize = 30, ^43^). The resulting 31 APPtg and 32 TAUtg modules were each summarized by the first principal component, known as module eigengenes (MEs). Next, Fisher’s Exact test with Benjamini-Yuketieli (BY) p-value adjustment was used to determine if a list of cell type-specific genes overlap significantly with genes in a module. More than half of all modules (APPtg: 16/31; TAUtg: 20/32) show significant overlap with a specific cell type (padj<0.05), particularly with the neuron and interneuron gene sets (APPtg: 10/31; TAUtg: 11/32; see Supplementary Fig.S3+4). Gene set enrichment analysis of GWAS genes from Marioni et al. ^2^ at different p-value cut-offs among the WGCNA-derived modules in APPtg or TAUtg mice was performed using Fisher’s exact test with Benjamini-Yuketieli (BY) p-value adjustment.

Functional annotations of the modules was performed using first GOrilla ^44^ and when no significant enrichment could be found, using DAVID (see Supplementary Tables 4 & 5.^45,46^ GO categories were deemed significant if the FDR-corrected p-value (GOrilla) or Benjamini-based p-value <0.05 (DAVID).

To search for potential regulators in each module, we ran i-cisTarget ^47^ which predicts transcription factor motifs (position weight matrices) and experimental data tracks (e.g. ENCODE, Roadmap Epigenomics Project) that are enriched in the input set of regions (i.e. genomic regions for each gene within the module). The default setting collected over 23588 features across all the databases available in i-cisTarget. Only regulators that were also expressed within the same WGCNA module were considered. Top regulators were selected based on the maximum normalized enrichment score for feature enrichment.

### Gene set overlap assessment

Overlap between the GWAS p<0.001 gene set (n=314), all genes significantly differentially expressed within the APPtg age*genotype interaction comparison (n=798), genes within the APPtg-Blue module (n=4236) and the microglia-specific gene set (n=435) were assessed using SuperExactTest^15^ (version 1.0.4), which calculates, based on combinatorial theory, the statistical probability of finding an over-representation of genes within the intersection of multiple sets, compared to random expectation. P-values were BY-adjusted (see Supplementary Table 7).

### Data and software availability

All data has been submitted to the GEO database. The mouse mRNAseq data is under accession number GSE110741. The software tools used for this study include: Tophat2 (version 2.0.7, ^36^), available from https://ccb.jhu.edu/software/tophat/index.shtml; Subread/Featurecounts ^48^ available from http://subread.sourceforge.net/; Pandas Python Data Analysis Library, available from http://pandas.pydata.org/; Limma/Linear Models for Microarray Data ^37^, available from https://bioconductor.org/packages/release/bioc/html/limma.html; Gene Set Enrichment Analysis ^40,41^, available from http://software.broadinstitute.org/gsea/index.jsp; WGCNA package in R ^42^, available from https://cran.r-project.org/web/packages/WGCNA/index.html; dynamicTreeCut package ^43^, available from https://cran.r-project.org/web/packages/dynamicTreeCut/index.html; Gene Ontology enrichment with GOrilla, ^44^ available from http://cbl-gorilla.cs.technion.ac.il/ and DAVID ^45,46^ available from https://david.ncifcrf.gov/home.jsp; i-cisTarget ^47^ available from https://gbiomed.kuleuven.be/apps/lcb/i-cisTarget/; SuperExactTest ^15^ available from https://cran.r-project.org/web/packages/SuperExactTest/index.html; StatsModels for Python available from http://www.statsmodels.org/0.8.0/generated/statsmodels.sandbox.stats.multicomp.multipletests.html.

### General statistical methods

In all instances of multiple testing Benjamini-Yuketieli p-value adjustment was performed in Python (http://www.statsmodels.org/0.8.0/generated/statsmodels.sandbox.stats.multicomp.multipletests.html) and α=0.05 throughout the study. When using Fisher’s exact test, one-sided p-value testing was performed; two-sided testing was performed in all other cases. More details for each individual statistical analysis can be found above.

## List of Supplementary Material

1. Supplementary Figures:
  1. Supplementary Fig.S1: Two APPtg-10M mice have lower transgene expression.
  2. Supplementary Fig.S2: APPtg and TAUtg mice demonstrate divergent neuronal and glial responses to increasing pathology load.
  3. Supplementary Fig.S3: Overview of APPtg-based WGCNA modules.
  4. Supplementary Fig.S4: Overview of TAUtg-based WGCNA modules.
  5. Supplementary Fig.S5: Examples of the mean z-score distribution of 10,000 randomly sampled gene sets with equal sizes to the gene set of interest.
2. Supplementary Materials & Methods
3. Supplementary Tables (separate Excel file, available upon request)
  1. Supplementary Table 1: Overview of the differential expression analysis per gene.
  2. Supplementary Table 2: Overview of the number of Marioni-based GWAS genes at different p-value cutoffs
  3. Supplementary Table 3: Overlap between genes within a WGCNA module and the cell-type specific gene sets.
  4. Supplementary Table 4: GOrilla-based GO enrichment per WGCNA module
  5. Supplementary Table 5: DAVID-based functional enrichment per WGCNA module
  6. Supplementary Table 6: Overview of all GWAS genes with p<0.5 that are significantly differentially expressed within the APPtg age*genotype interaction comparison.
  7. Supplementary Table 7: Results from the SuperExactTest for overlaps between multiple gene sets.

## Acknowledgements

We thank Veronique Hendrickx and Jonas Verwaeren for animal husbandry, and Carlo Sala Frigerio and Tom Jaspers for discussions. APPtg mice were a kind gift from Mathias Jucker, DZNE, Germany.

## Funding

Annerieke Sierksma and Bart De Strooper are supported by the Opening the Future campaign of the Leuven Universitair Fonds (LUF) and the Alzheimer Research Foundation (SAO-FRA; P#16017). Evgenia Salta is an FWO (12A5316N) and Alzheimer’s Association (AARF-16-442853) postdoctoral fellow. Work in the De Strooper laboratory was supported by the Fonds voor Wetenschappelijk Onderzoek (FWO), KU Leuven, VIB, and a Methusalem grant from KU Leuven and the Flemish Government, Vlaams Initiatief voor Netwerken voor Dementie Onderzoek (VIND, Strategic Basic Research Grant 135043) and the “Geneeskundige Stichting Koningin Elisabeth”. Bart De Strooper is holder of the Bax-Vanluffelen Chair for Alzheimer’s Disease. Luc Buée is supported by grant ANR-16-COEN-0007.

## Author Contributions

A.S., B.D.S. and M.F. designed the study and wrote the manuscript. A.S., A.L., J.Z. and M.F. performed analyses. D.B. and L.B. provided TAUtg mice. E.S., R.M., D.B. and L.B. provided expertise and feedback. All authors read and approved the final manuscript for publication.

## Competing interests

B.D.S. is ad hoc consultant for various companies but has no direct financial interest in the current study. L.B. is consultant for Servier and Remynd. He receives research funding from UCB Pharma, but not for the work presented in the current manuscript. A.S., A.L., E.S., R.M., J.Z., D.B. and M.F. report no biomedical financial interests or potential conflicts of interest.

## Data availability

The mouse mRNAseq data has been submitted to the GEO database under accession number GSE110741. All other data are available in the main text or the supplementary materials and tables.

