## Supplementary for "Novel Alzheimer risk genes determine the microglia response to amyloid-β but not to TAU pathology"

### Supplementary Information to Sierksma et al.

#### Overview of Supplementary Material

1. Supplementary Figures:
    1. Supplementary Fig.S1: Two 10M APPtg mice have lower transgene expression.
    2. Supplementary Fig.S2: APPtg and TAUtg mice demonstrate divergent neuronal and glial responses to increasing pathology load.
    3. Supplementary Fig.S3: Overview of APPtg-based WGCNA modules.
    4. Supplementary Fig.S4: Overview of TAUtg-based WGCNA modules.
    5. Supplementary Fig.S5: Examples of the mean z-score distribution of 10,000 randomly sampled gene sets with equal sizes to the gene set of interest.
2. Supplementary Materials & Methods
3. Supplementary Tables (separate Excel file)
    1. Supplementary Table 1: Overview of the differential expression analysis per gene.
    2. Supplementary Table 2: Overview of the number of Marioni-based GWAS genes at different p-value cutoffs
    3. Supplementary Table 3: Overlap between genes within a WGCNA module and the cell-type specific gene sets.
    4. Supplementary Table 4: GOrilla-based GO enrichment per WGCNA module
    5. Supplementary Table 5: DAVID-based functional enrichment per WGCNA module
    6. Supplementary Table 6: Overview of all GWAS genes with p<0.5 that are significantly differentially expressed within the APPtg age*genotype interaction comparison.
    7. Supplementary Table 7: Results from the SuperExactTest for overlaps between multiple gene sets.

### Supplementary Figures and Figure Legends

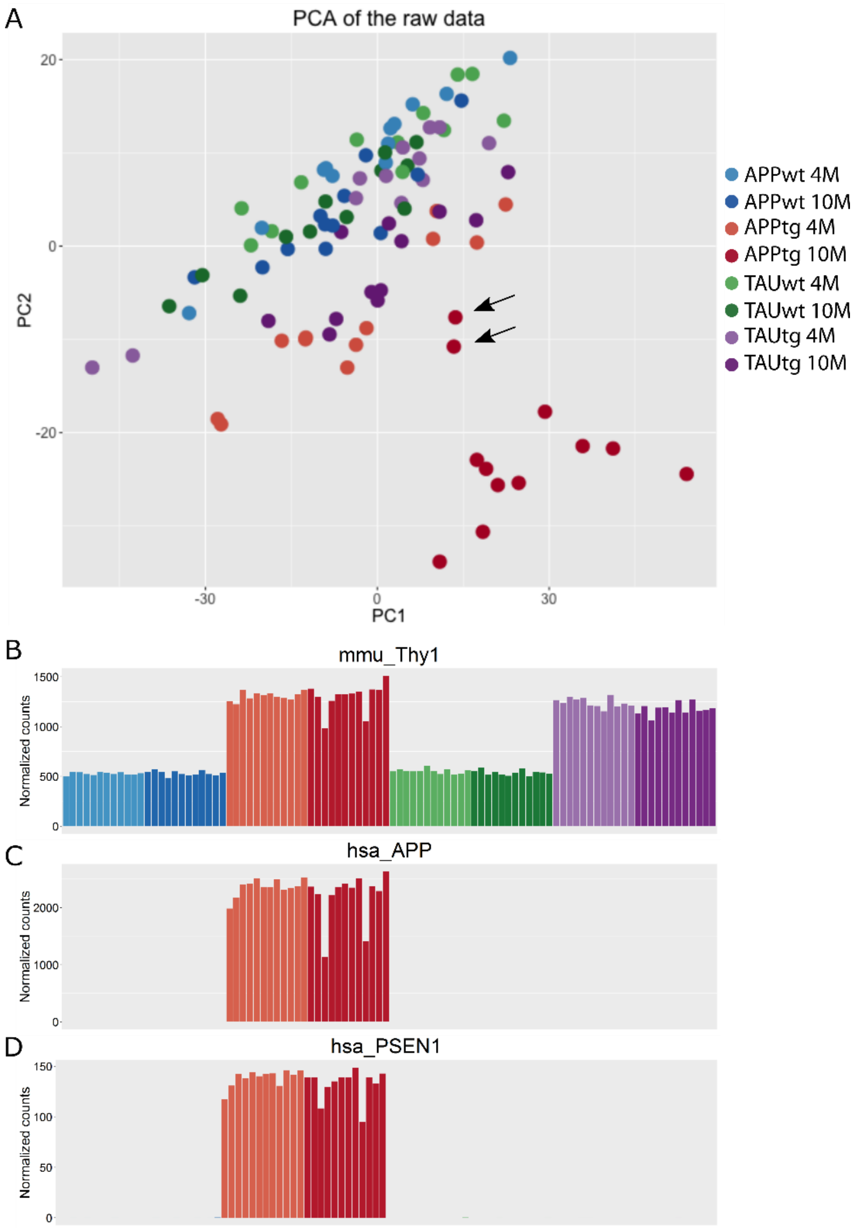


Supplementary Fig.S1: Two APPtg-10M mice have lower transgene expression. A: The first 2 principle components of the PCA plot of the log2(raw reads from Feature Counts +1) demonstrate that 2 APPtg-10M mice cluster more towards the APPtg-4M mice (see arrows). B-D: These two mice have decreased expression of transgenes mmu-Thy1 (-24%; B), hsa-APP (-46%; C) and hsa-PSEN1 (-26%; D) compared to other APPtg-10M mice. Expression of hsa-MAPT, mmu-App, mmu-Psen1 and mmu-Mapt remained unaffected (data not shown).

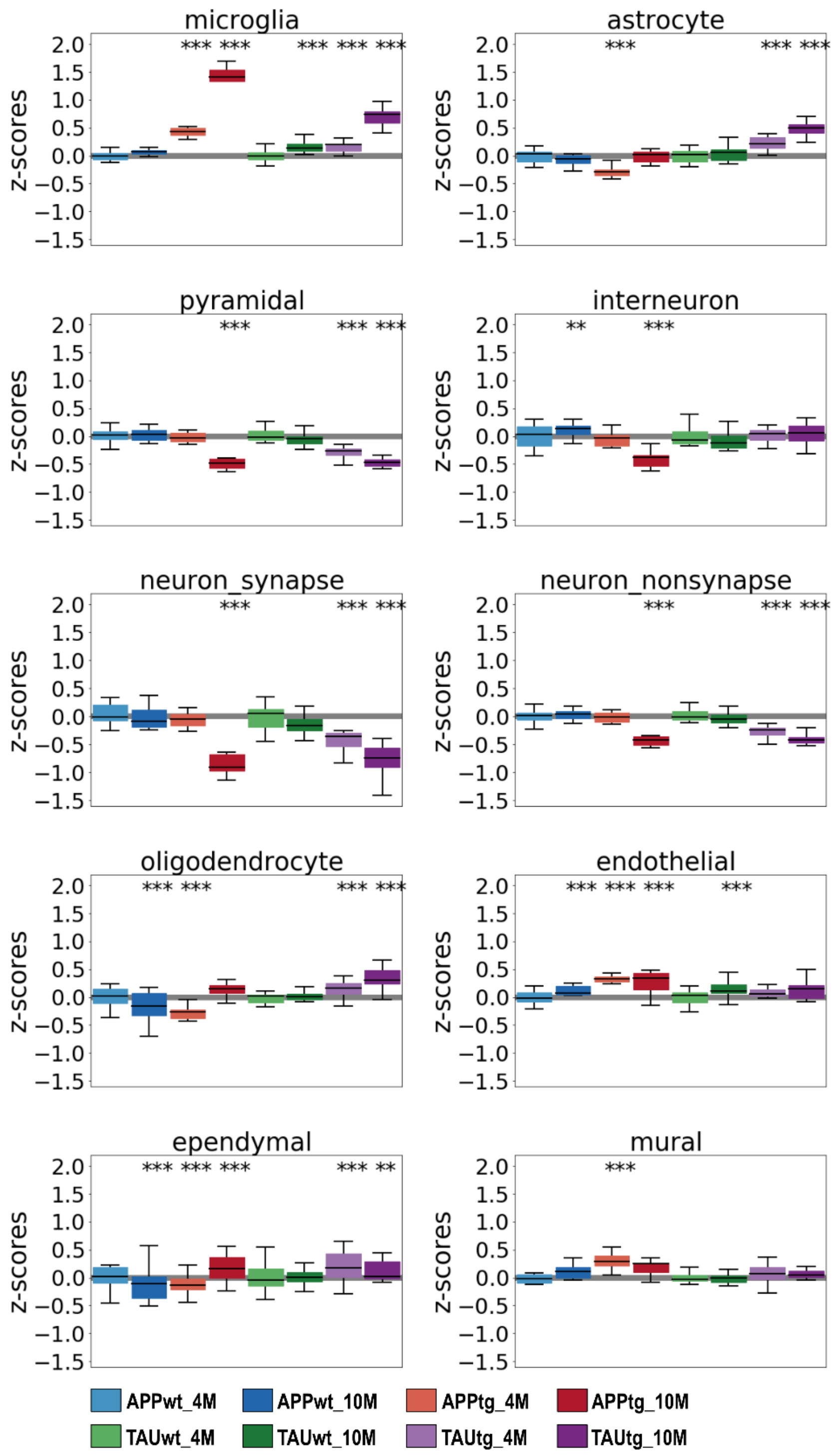


**Supplementary Fig.S2: APPtg and TAUtg mice demonstrate divergent neuronal and glial responses to increasing pathology load.** Z-score distribution per experimental group for all genes in cell type specific gene sets for as defined by Zeisel et al. *(17)* and SynaptomeDB *(47)*. Empirical p-values are

Bonferroni adjusted (pbonf) and indicate significant shift in z-score distribution (see Supplementary Materials & Methods). ***: pbonf <0.001, **: pbonf 0.01.

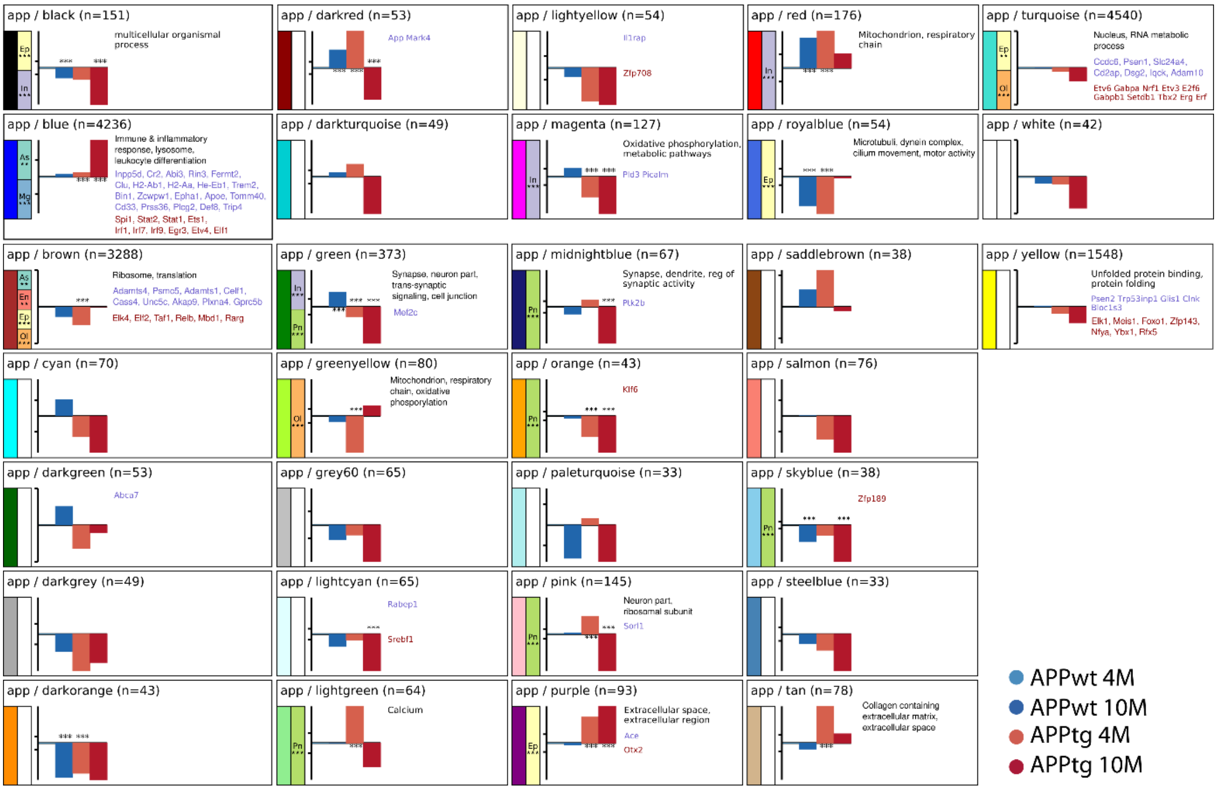


**Supplementary Fig.S3: Overview of APPtg-based WGCNA modules.** For each module is indicated the z-score distribution of each experimental group and their bonferroni-adjusted empirically-derived p-values. If significant GO enrichment is observed, the categories are mentioned in the top row. If the module contains established GWAS genes, they are reported on the middle row. If there is an enrichment of transcription factor targets of transcription factors expressed in that module, the transcription factors are labeled on the bottom row. If significant enrichment with a specific cell type gene set is observed, this is indicated with the cell-type labeled colored boxes. As: astrocytes; En: Endothelial; Ep: Ependymal; In: interneurons; Mg: microglia; Ol: oligodendrocyte; Pn: Pyramidal neuron.

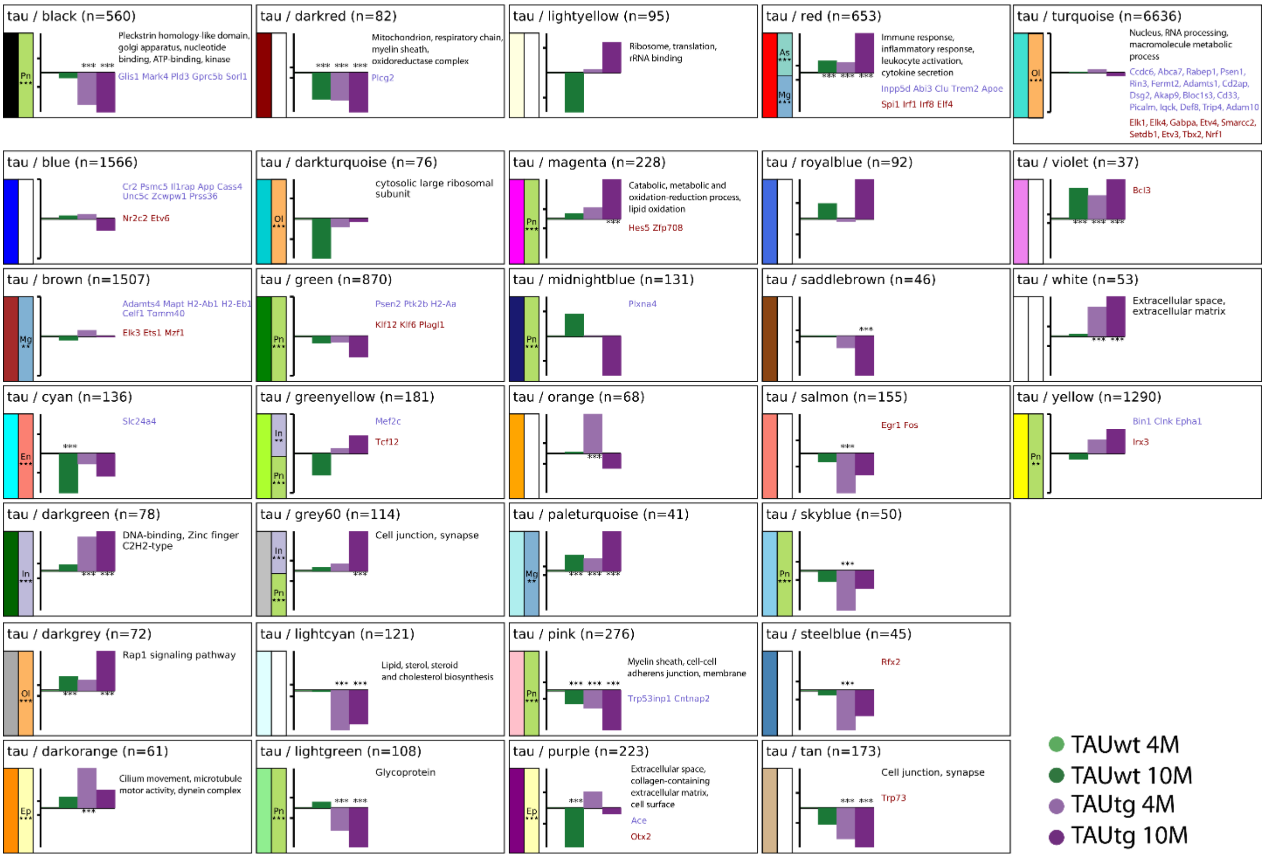


**Supplementary Fig.S4: Overview of TAUtg-based WGCNA modules.** For each module is indicated the z-score distribution of each experimental group and their bonferroni-adjusted empirically-derived p-values. If significant GO enrichment is observed, the categories are mentioned in the top row. If the module contains established GWAS genes, they are reported on the middle row. If there is an enrichment of transcription factor targets of transcription factors expressed in that module, the transcription factors are labeled on the bottom row. If significant enrichment with a specific cell type gene set is observed, this is indicated with the cell-type labeled colored boxes. As: astrocytes; En: Endothelial; Ep: Ependymal; In: interneurons; Mg: microglia; Ol: oligodendrocyte; Pn: Pyramidal neuron.

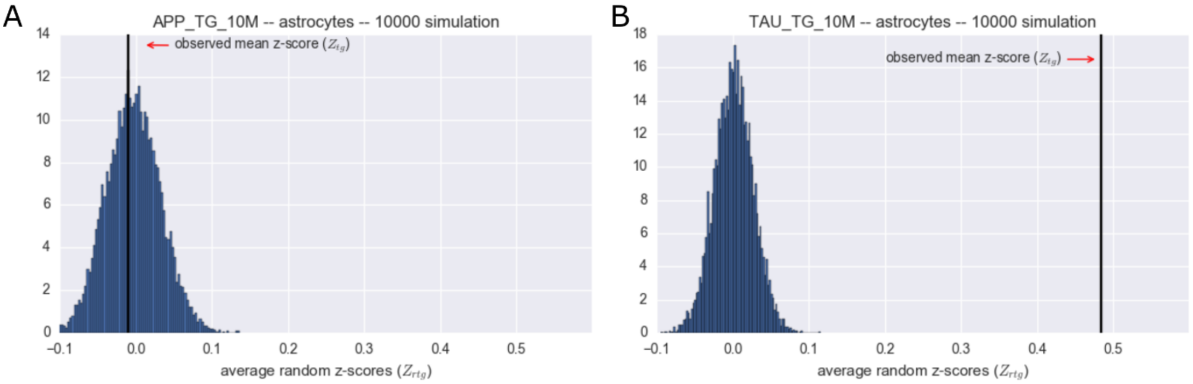


**Supplementary Fig.S5: Examples of the mean z-score distribution of 10,000 randomly sampled gene sets with equal sizes to the gene set of interest (e.g. 719 astrocyte genes).** The black vertical line represents the observed mean z-score $Z_{tg}$ for APP-TG-10M (A) or TAU-TG-10M mice (B). Its cumulated distribution function among the normally distributed random z-scores $Z_{rtg}$ determines the associated empirical p-value.

### Supplementary Materials & Methods

#### Empirical p-values associated with predicted changes in cell type fraction

We predict changes in cell type fractions based on shifts in the mean (per gene z-normalized across samples) expression of a wide panel of genes associated to a specific cell type by Zeisel et al. *(17)*. Given that we are interested in shifts of cell types when compared to the 4M old wild-type (WT-4M) mice (from the corresponding strain) we use the mean gene expression in only the WT-4M mice as the baseline in the gene expression z-normalization. So, the z-normalized expression $Z_i$ of a gene *i* in sample *s* in can be calculated from the (limma-voom normalized) expression $E_i$ as follows:

$$Z_{is} = \frac{E_{is} - mean\left(E_{i(WT.4M)}\right)}{std(E_i)}$$

Then the average z-score ($Z_{tg}$) for a cell type (*t*), in an experimental group (*g*) across all genes of a cell type gene set is calculated as the mean of the *per gene* z scores ($Z_{is}$) in the experimental group. It is these $Z_{tg}$ scores that are depicted in Fig. 3A and Supplementary Fig.S3.

Next, an empirical p-value is assigned to each $Z_{tg}$ as follows: per cell type specific gene set, 10,000 random gene sets (of the same size as the cell type specific gene set) were sampled. For each random iteration a $Z_{rtg}$ is calculated using the z-normalized $Z_{is}$ scores in exactly the same way as described above. Of each population of randomized $Z_{rtg}$ values (per group *g* and celltype *t*), normality is confirmed and a population mean and standard deviation are calculated to assign an empirical p-value to the observed $Z_{tg}$ using a cumulative distribution function (Supplementary Fig.S5). The combined observations across 6 experimental groups (excluding both WT-4M) and 6 different cell gene sets types are Bonferroni corrected to exclude false-positive observations.
